# Experiments and simulations on short chain fatty acid production in a colonic bacterial community

**DOI:** 10.1101/444760

**Authors:** Bea Yu, Ilija Dukovski, David Kong, Johanna Bobrow, Alla Ostrinskaya, Daniel Segrè, Todd Thorsen

**Affiliations:** MIT Lincoln Lab, 244 Wood Street, Lexington, MA 02421, USA; Bioinformatics Graduate Program, Boston University, Boston, MA 02215, USA; Biological Design Center, Boston University, Boston, MA 02215, USA; Director, Community Biotechnology Initiative, MIT Media Lab, Massachusetts Institute of Technology, 77 Mass. Ave., E14/E15, Cambridge, MA 02139 USA; Department of Biology, Department of Biomedical Engineering and Department of Physics, Boston University, Boston, MA 02215, USA

## Abstract

Understanding how production of specific metabolites by gut microbes is modulated by interactions with surrounding species and by environmental nutrient availability is an important open challenge in microbiome research. As part of this endeavor, this work explores interactions between *F. prausnitzii,* a major butyrate producer, and *B. thetaiotaomicron*, an acetate producer, under three different *in vitro* media conditions in monoculture and coculture. *In silico* Genome-scale dynamic flux balance analysis (dFBA) models of metabolism in the system using COMETS (Computation of Microbial Ecosystems in Time and Space) are also tested for explanatory, predictive and inferential power. Experimental findings indicate enhancement of butyrate production in coculture relative to *F. prausnitzii* monoculture but defy a simple model of monotonic increases in butyrate production as a function of acetate availability in the medium. Simulations recapitulate biomass production curves for monocultures and accurately predict the growth curve of coculture total biomass, using parameters learned from monocultures, suggesting that the model captures some aspects of how the two bacteria interact. However, a comparison of data and simulations for environmental acetate and butyrate changes suggest that the organisms adopt one of many possible metabolic strategies equivalent in terms of growth efficiency. Furthermore, the model seems not to capture subsequent shifts in metabolic activities observed experimentally under low-nutrient regimes. Some discrepancies can be explained by the multiplicity of possible fermentative states for *F. prausnitzii*. In general, these results provide valuable guidelines for design of future experiments aimed at better determining the mechanisms leading to enhanced butyrate in this ecosystem.

**Importance:** Studies associating butyrate levels with human colonic health have inspired research on therapeutic microbiota consortia that would optimize butyrate production if implanted in the human colon. *Faecalibacterium prausnitzii* is commonly observed in human fecal samples and produces butyrate as a product of fermentation. Previous studies indicate that *Bacteroides thetaiotaomicron*, also commonly found in human fecal samples, may enhance butyrate production in *F. prausnitzi* when the two species are co-localized. This possibility is investigated here under different environmental conditions using experimental methods paired with computer simulations of the whole metabolism of bacterial cells. Initial findings indicate that interactions between these two species result in enhanced butyrate production. However, results also paint a nuanced picture, suggesting the existence of a multiplicity of equivalently efficient metabolic strategies and complex interactions between acetate and butyrate production in these species that appear highly dependent on specific environmental conditions.

## Introduction

It is increasingly recognized that metabolites produced by the resident microbiota of the colon have a major influence on host physiology (1). Dietary substrates dramatically influence the amount and type of these metabolites produced (2). For instance, fermentation of carbohydrates produces a number of bioactive compounds, most notably short chain fatty acids (SCFA) such as butyrate, that have been demonstrated to shape the gut microenvironment, serve as an energy source for the colonic epithelium, and influence disease through anti-inflammatory, lipogenic, and anti-apoptotic effects (3–6).

The production of metabolites in a microbial community has been suggested to be heavily modulated by interactions among its members. These interactions manifest in a variety of modes, ranging from competitive or predatory to commensal and mutualistic exchanges (7). Additionally, many microbes in nature exist in spatially defined structures (8), such as the mucosal layer of the gut. Spatial assortment of cells creates locally heterogeneous subpopulations with varying access to resources that can also modulate inter- and intra-community behavior (9). A major goal of ongoing efforts in human microbiome research (10) is to gain enough predictive and quantitative understanding of inter-microbial interactions (11) and of the metabolic interplay between microbiota and host (12) to be able to understand the effects of the microbiome on human health. These capabilities could greatly facilitate successful design of therapeutic strategies for microbiome-related diseases.

Efforts towards safely, effectively and reliably engineering microbial communities (9) to improve human health are, however, limited by insufficient understanding of the nature of the mechanisms underlying microbial interactions and the way these interactions affect microbiome dynamics. Anaerobic *in vitro*, *in vivo* and *ex vivo* experiments capable of probing systems similar to the human colonic environment are difficult and expensive. Previous work uncovering fundamental properties of SCFA-producing bacteria and their symbiotic partners has used *Faecalibacterium prausnitzii* as a model system (13–21). This is motivated by the high prevalence of *F. prausnitzii* as a commensal bacterium in the human large intestine (22) and the role it plays as one of the major butyrate producers (23). Among the bacteria used for coculture studies, a common gut commensal, *Bacteroides thetaiotaomicron,* has been chosen in both experimental (24) and computational (25) analyses of SCFA production. In particular, experimental efforts to grow *F. prausnitzii* in coculture with *B. thetaiotaomicron* have suggested enhancement of butyrate production in coculture relative to *F. prausnitzii* monoculture (24). However, the data in this study was obtained only for a single time point and no information on the dynamics of the biomass or butyrate production was provided. In general, to our knowledge, no comparison has been previously made between experimentally measured time-courses of the biomass of these species and their respective metabolic dynamics, when grown individually and in co-culture.

In parallel, computational work based on metabolic network analyses has led to the construction of genome scale models for each of these bacterial species, and to a computational assessment of their metabolic capabilities (26–28). The modeling approach used in these studies, often referred to as constraint-based modeling (or stoichiometric modeling) is based on simplifying assumptions about the intracellular dynamics of metabolism. It enables quantitative predictions of the intracellular and exchange fluxes, in addition to the growth rate of different species. In particular, flux balance analysis (FBA) (29), can be used to calculate the flow of metabolites through a metabolic network, making it possible to predict the growth rate of an organism or the rate of production of important metabolites (30) (see also (31) for a comprehensive review of different approaches). While the reconstructed networks and the modeling tools used for making these predictions vary widely in accuracy and predictive power, the formal representation of metabolism into these mathematical structures and codification of multi-level processes into algorithms have sparked a revolution in systems biology of metabolism, enabling precise hypothesis testing, and the formulation of genome-scale based community modeling. In the context of human gut microbiome studies and inter-species interactions, modeling work has been shown in particular to provide insight into the stability of biofilm forming communities (25).

In order to make comparisons between computational predictions and experimental time-course data, it is important to be able to connect detailed knowledge of the intracellular metabolism of individual organisms to the dynamic metabolic changes occurring in the surrounding environment. An extension of FBA capable of these types of calculations is dynamic FBA (or dFBA)(32). Harcombe *et al.* (33) developed a computational framework specifically designed to help predict the spatio-temporal behavior of synthetic microbial consortia. This system, known as Computation of Microbial Ecosystems in Time and Space (COMETS) (33), generates predictions of biomass growth curves as well as detailed time dynamics of the concentrations of all nutrients and metabolites in the environment. COMETS has been shown to accurately predict the behavior of small artificial ecosystems.

Despite the availability of these experimental data and computational tools, many fundamental features of the interactions between *F. prausnitzii* and *B. thetaiotaomicron*, as well as our capacity to predict clinically relevant variables, remain unexplored. While previous studies have computationally and experimentally analyzed the metabolic capabilities of each of these bacteria individually (26, 27) and the dependence of these and other bacteria upon different oxygen levels (25), no direct comparison of experimental and computational time course data for this consortium under varying conditions has been presented before. In particular, no attempt has been made to recapitulate or predict these time-courses with dynamic computational models.

Here, we provide novel insight into the *F. prausnitzii* - *B. thetaiotaomicron* model system by combining new experimental measurements of bacterial biomass and environmental metabolites with COMETS-based computer simulations. We performed a series of anaerobic *in vitro* experiments involving monocultures and cocultures of *F. prausnitzii* and *B. thetaiotaomicron* grown in three different media, and found increased butyrate production in co-culture relative to monoculture under high glucose and acetate concentrations. Upon fitting of six parameters for metabolic uptake kinetics in monocolture, COMETS simulations were able to recapitulate biomass time courses in monoculture and predict combined biomass time courses for coculture. Model predictions for butyrate, however, portray a more complex picture. Accurate predictions of initial butyrate production rate do not hold at longer times due to the existence of multiple alternative optima in the flux states and the history-dependence of the dynamical predictions. Strong sensitivity of the butyrate production curves to specific concentrations of nutrients, including phosphate, provide insight into the complexity of these metabolic exchanges, and valuable guidance for future experimental and modeling work.

## Results

### *In vitro* and *in silico* coculture biomass dynamics under different nutrient limitations

We initially characterized anaerobic growth of *B. thetaiotaomicron* and *F. prausnitzii* individually and in coculture, under different levels of carbon availability (low, medium, high, see Methods). In addition to glucose, acetate was added proportionally, mimicking the fermentative activity of the rest of the microbiota (13). The presence of acetate in the medium also allowed us to assess how, even in the absence of *B. thetaiotaomicron, F. prausnitzii* responds to varying acetate availability.

Monoculture growth for *B. thetaiotaomicron* appears sensitive to the amount of carbon provided (Fig. 1). Growth rate and yield increase in medium acetate/glucose medium compared to low acetate/glucose concentrations. No clear increase in biomass occurred when carbon abundance was increased from medium to high levels. The amount of *F. prausnitzii* biomass in monoculture appears insensitive to increase in initial acetate/glucose levels. The combined biomass growth curves of the coculture (measured as a collective OD) closely tracks OD curves of *B. thetaiotaomicron*, suggesting a prominent role of this bacterium in the consortium. This observation is consistent with previous experiments (3) in which the combined coculture biomass OD of *B. adolescentis* and *F. prausnitzii* mirrors the OD of the *B. adolescentis* monoculture, suggesting that some features of that consortium may be similar to the one studied here, even across different spatial scales and environmental settings.

**Figure 1.**
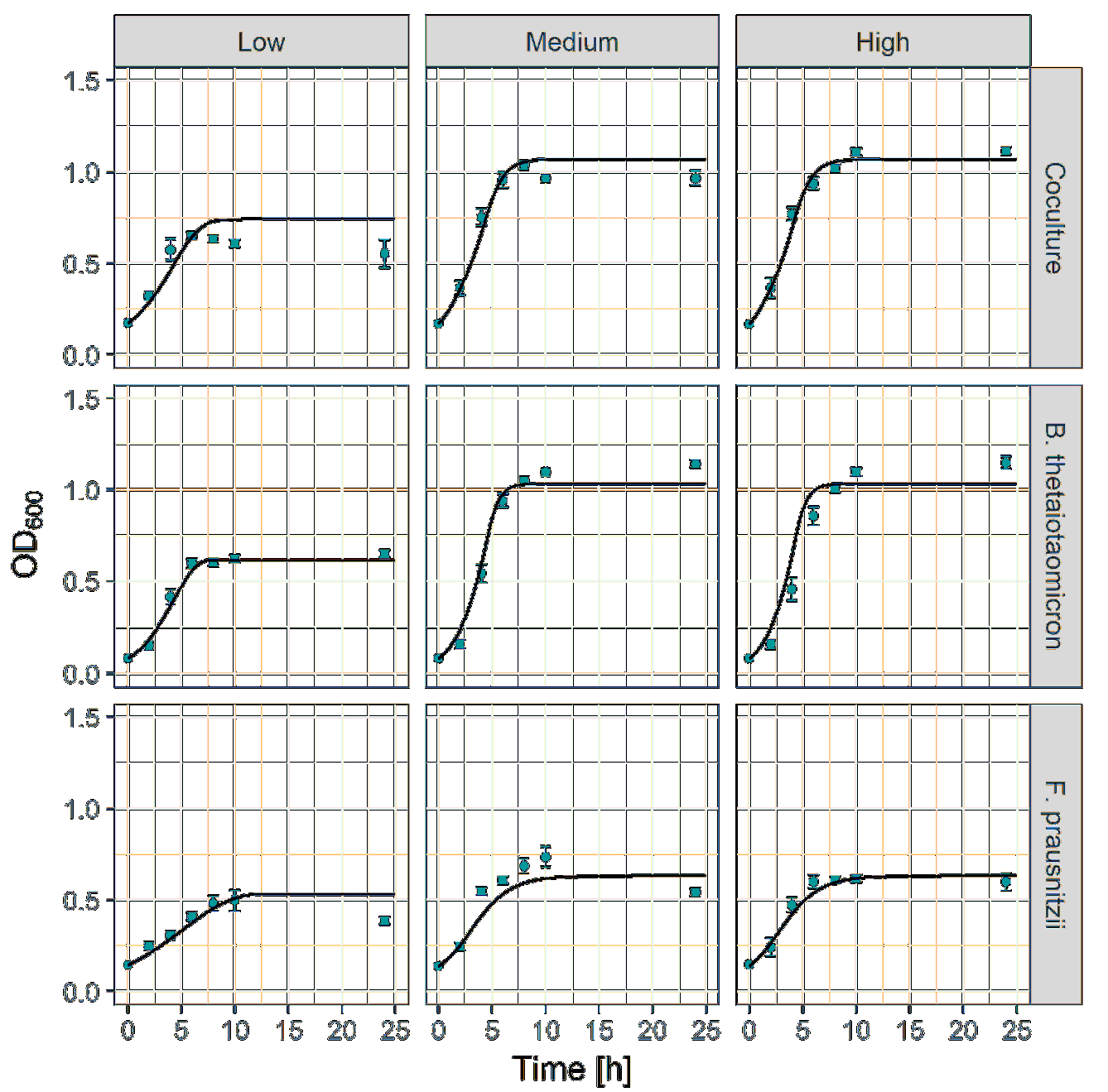
Optical densities for single species and coculture of *F. prausnitzii* and *B. thetaiotaomicron*, grown in three different media conditions. The simulations (solid curves) were obtained by dFBA with the same set of uptake parameters for all media conditions. The columns correspond to the media conditions while the rows correspond to the cultured species.

In parallel to the experimental measurements, we implemented *in silico* simulations of the same monocultures and cocultures, using previously published genome-scale metabolic models for the two bacteria (26, 27). In particular, we used COMETS (33) to test whether (i) parameters from the literature, combined with minimal fitting of unknown parameters, would recapitulate the observed monoculture behavior and (ii) models tuned for monoculture experiments would be adequate to predict the outcome of the coculture experiment.

Superimposed on the experimental data, Fig. 1 shows the biomass dynamics as simulated in COMETS. The monoculture simulations were supplemented with empirical knowledge of uptake *K_M_* and fitting of *V_max_* values. After selecting initial kinetic parameter values based on previously determined corresponding parameters for the phosphotransferase (PTS) transporter (21, 34), a sensitivity analysis allowed us to identify *V_max_* values that provide best fit of growth curves to monoculture (Fig. 2). This calibration step, similar to that previously performed in (35), produces simulated OD curves that broadly agree with the experimentally measured points (see root-mean-squared error (RMSE) in Table 1). Using these parameters for COMETS simulations of cocultures at the experimentally estimated initial biomass abundances, coculture predictions track coculture experimental data closely, as shown by predictive RMSE values (Table 1).

**Figure 2.**
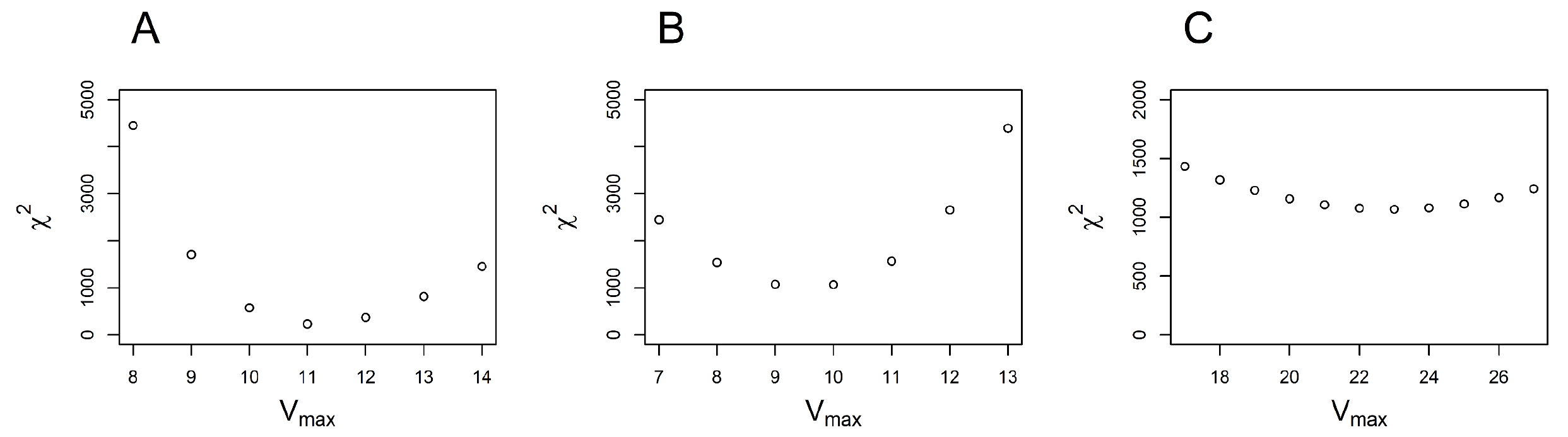
The sensitivity of the OD curves to the values of the maximum nutrient uptake parameter *V_max_*. The values of the *V_max_* parameter used in the simulations were obtained by minimizing χ^2^,the sum of the squared deviations of the simulation from the experimental values, weighted by the measured variance. We used a single value of *V_max_* for the uptake of all nutrients by the *B. thetaiotaomicron* model, with the minimum of minimizing χ^2^, shown in panel A). In the case of *F. prausnitzii*, we determined two separate values of minimizing χ^2^, one for glucose and acetate uptake shown on panel B), and another one for the rest of the nutrients, shown on panel C).

**Table 1.**
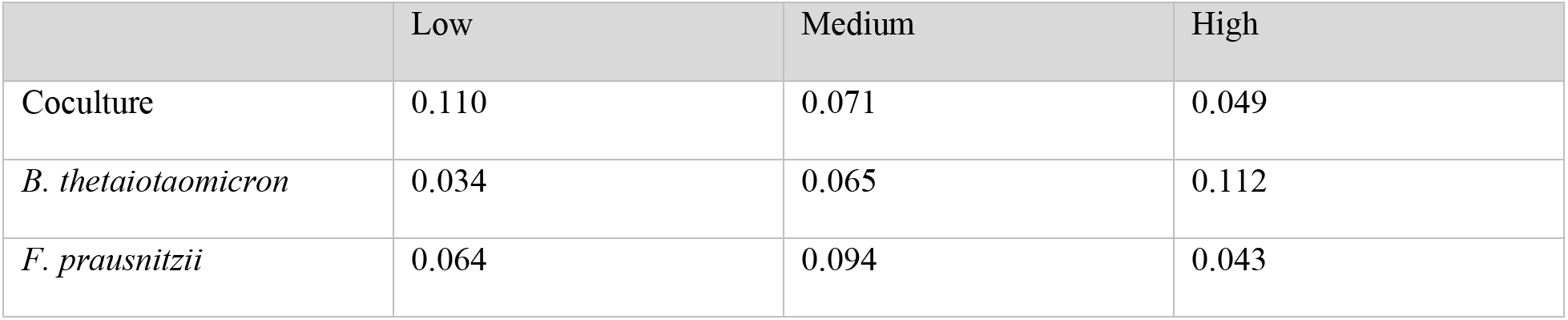
Table of RMSE between the values in Fig. 1 measured experimentally and predicted by simulations, for monocultures and coculture in three carbon source conditions as described in the text.

### Coculture conditions impact the average rate of butyrate production in experiments

Throughout all of our experiments, in addition to monitoring the overall OD, we measured the extracellular aboundance of butyrate and acetate. In Fig. 3, the average amount of butyrate produced by *F. prausnitzii* in coculture appears higher than monoculture in the medium and high initial acetate/glucose concentrations, but not in the low concentration conditions. In low acetate/glucose conditions the butyrate production rate in coculture appears suppressed relative to the one in monoculture. Box’s approach in (36), applying ANOVA to summary statistics describing growth curves, was used to quantify the statistical significance of the difference in butyrate production curves across different treatments, i.e.: (i) monoculture vs. coculture conditions (mono/co) and (ii) the three different initial acetate/glucose initial concentrations (initial glu/ac). Tables 2 and 3 show that the average rate of butyrate production in *F. prausnitzii* is significantly altered in mono vs. coculture conditions but not across the different initial abundances of acetate and glucose. The initial acetate/glucose concentrations, but not mono vs. coculture conditions, significantly change the average rate of acetate production from *B. thetaiotaomicron*.

**Figure 3.**
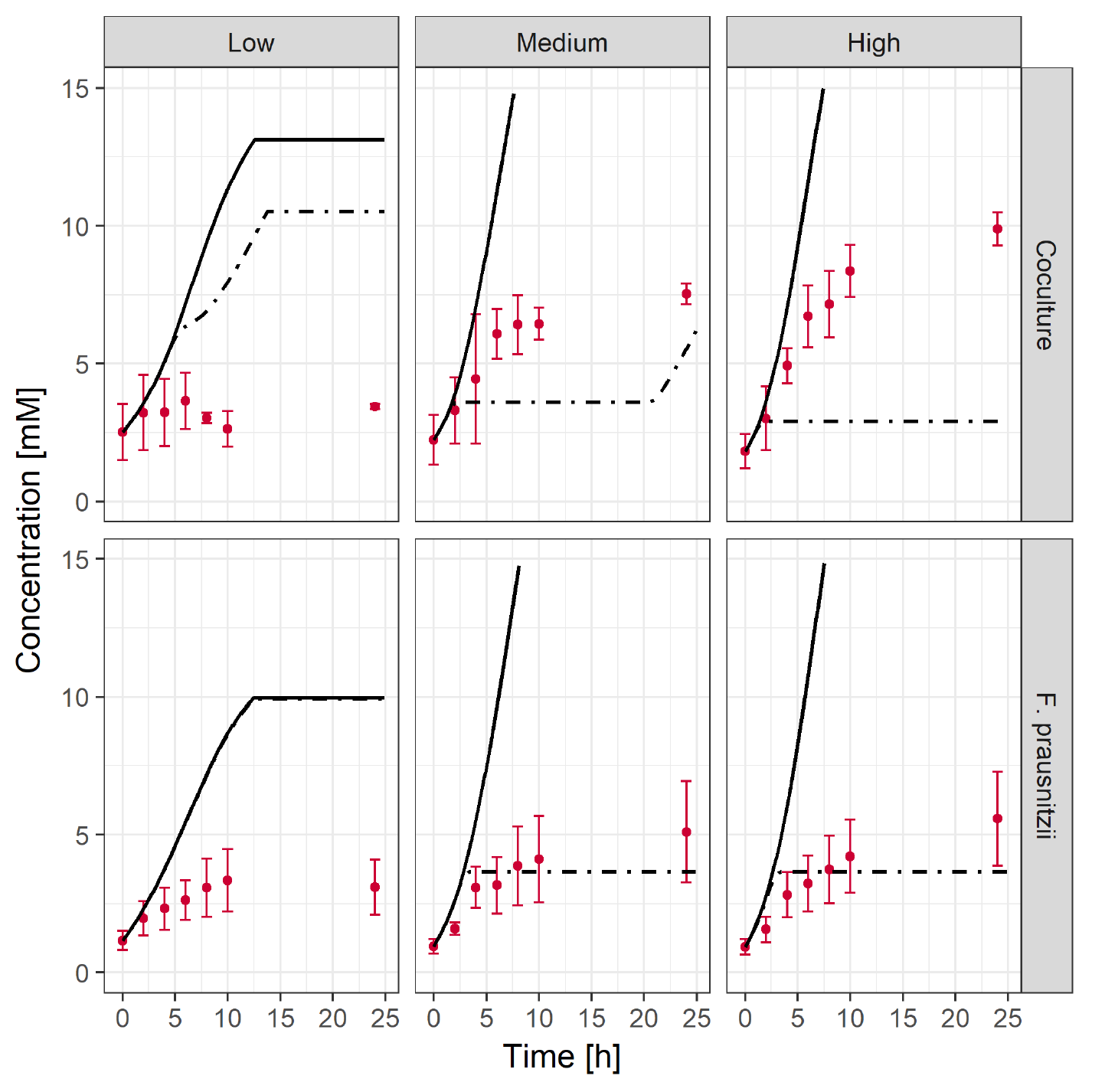
Experimental and simulated (solid, dot and dash curves) butyrate production time courses for monocultures and coculture under the three initial glucose/acetate initial concentrations. Apparent differences demonstrated in these plots in butyrate production for *F. prausnitzii* monoculture vs. coculture and for the three initial concentrations can be tested statistically using ANOVA on summary statistics describing the curves. The simulated curves correspond to the maximized (solid curve) and minimized (dash dot curve) butyrate secretion.

**Table 2.** Analysis of Variance of Shape of Metabolite Curves.

|  | Acetate |  |  |  |  | Butyrate |  |  |  |  |
| --- | --- | --- | --- | --- | --- | --- | --- | --- | --- | --- |
| Source | Sum Sq. | d.f. | Mean Sq. | F | Prob>F | Sum Sq. | d.f. | Mean Sq. | F | Prob>F |
| initial glc/ac | 21.14 | 2 | 10.571 | 0.47 | 0.6277 | 1.3206 | 2 | 0.66032 | 0.69 | 0.5015 |
| mono/co | 115.13 | 1 | 115.131 | 5.1 | 0.0261 | 0.11956 | 1 | 0.11945 | 0.13 | 0.7237 |
| initialglc/ac*mono/co | 76.86 | 2 | 38.43 | 1.7 | 0.1877 | 0.275 | 2 | 0.13751 | 0.14 | 0.8655 |
| Error | 2304.56 | 102 | 22.594 |  |  | 96.9437 | 102 | 0.95043 |  |  |
| Total | 2517.69 | 107 |  |  |  | 98.6588 | 107 |  |  |  |

**Table 3.**
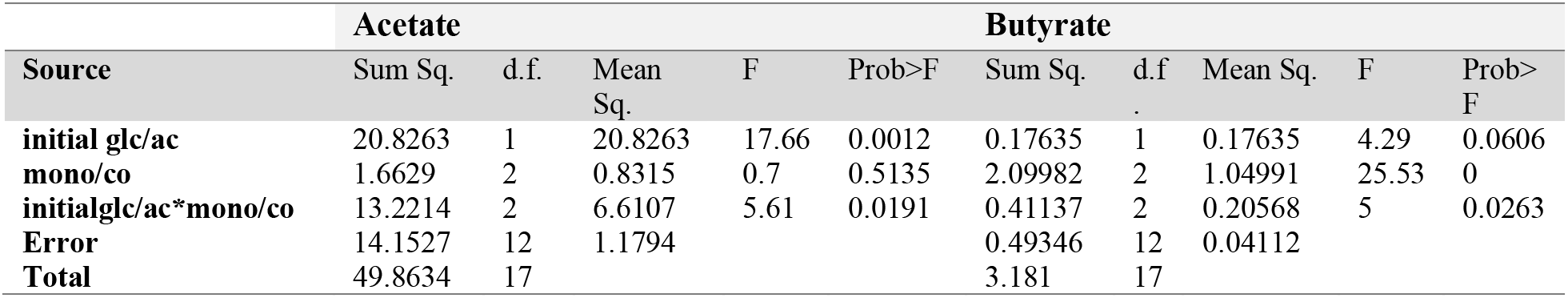
Analysis of Variance of Average Rate of Metabolite Curves.

|  | Acetate |  |  |  |  | Butyrate |  |  |  |  |
| --- | --- | --- | --- | --- | --- | --- | --- | --- | --- | --- |
| Source | Sum Sq. | d.f. | Mean Sq. | F | Prob>F | Sum Sq. | d.f. | Mean Sq. | F | Prob>F |
| initial glc/ac | 20.8263 | 1 | 20.8263 | 17.66 | 0.0012 | 0.17635 | 1 | 0.17635 | 4.29 | 0.0606 |
| mono/co | 1.6629 | 2 | 0.8315 | 0.7 | 0.5135 | 2.09982 | 2 | 1.04991 | 25.53 | 0 |
| initialglc/ac*mono/co | 13.2214 | 2 | 6.6107 | 5.61 | 0.0191 | 0.41137 | 2 | 0.20568 | 5 | 0.0263 |
| Error | 14.1527 | 12 | 1.1794 |  |  | 0.49346 | 12 | 0.04112 |  |  |
| Total | 49.8634 | 17 |  |  |  | 3.181 | 17 |  |  |  |

**Table 4.**
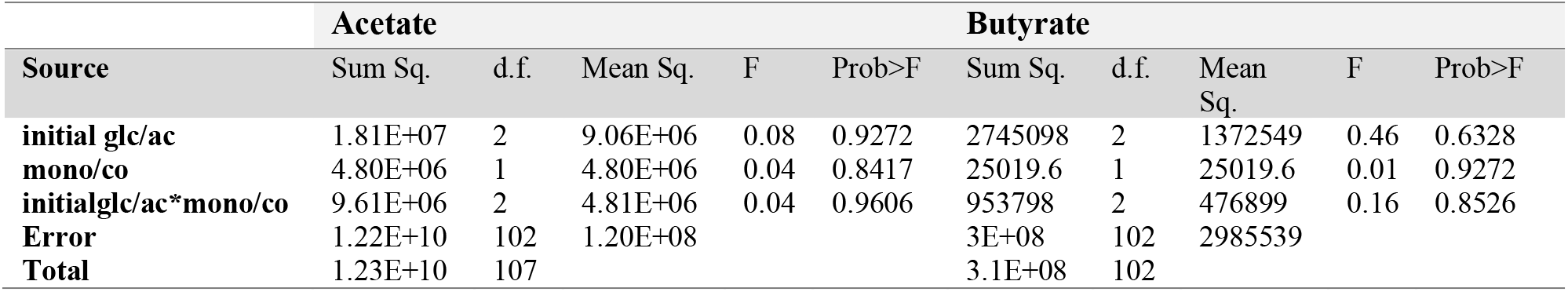
Analysis of Variance of Average Rate of Metabolite Curves.

|  | Acetate |  |  |  |  | Butyrate |  |  |  |  |
| --- | --- | --- | --- | --- | --- | --- | --- | --- | --- | --- |
| Source | Sum Sq. | d.f. | Mean Sq. | F | Prob>F | Sum Sq. | d.f. | Mean Sq. | F | Prob>F |
| initial glc/ac | 1.81E+07 | 2 | 9.06E+06 | 0.08 | 0.9272 | 2745098 | 2 | 1372549 | 0.46 | 0.6328 |
| mono/co | 4.80E+06 | 1 | 4.80E+06 | 0.04 | 0.8417 | 25019.6 | 1 | 25019.6 | 0.01 | 0.9272 |
| initialglc/ac*mono/co | 9.61E+06 | 2 | 4.81E+06 | 0.04 | 0.9606 | 953798 | 2 | 476899 | 0.16 | 0.8526 |
| Error | 1.22E+10 | 102 | 1.20E+08 |  |  | 3E+08 | 102 | 2985539 |  |  |
| Total | 1.23E+10 | 107 |  |  |  | 3.1E+08 | 102 |  |  |  |

Interpretation of the above assessments of butyrate and acetate production is limited by the lack of experimental knowledge of the precise amount of biomass of each species in the coculture experiments. In particular, in absence of further laborious organism-specific data, it is impossible to determine whether significant changes in the level of the butyrate curves are due to *F. prausnitzii* producing more butyrate at the cellular level in the coculture, or whether the increase is due to an increase in *F. prausnitzii* total biomass, enabled by coculture conditions, or both. Under these circumstances, COMETS-predicted biomass estimates for each species from coculture simulations (Fig. 6B) enabled hybrid computational-experimental estimates of biomass-normalized butyrate (Fig. 6A) and acetate (Fig. S2) production curves.

**Figure 4.**
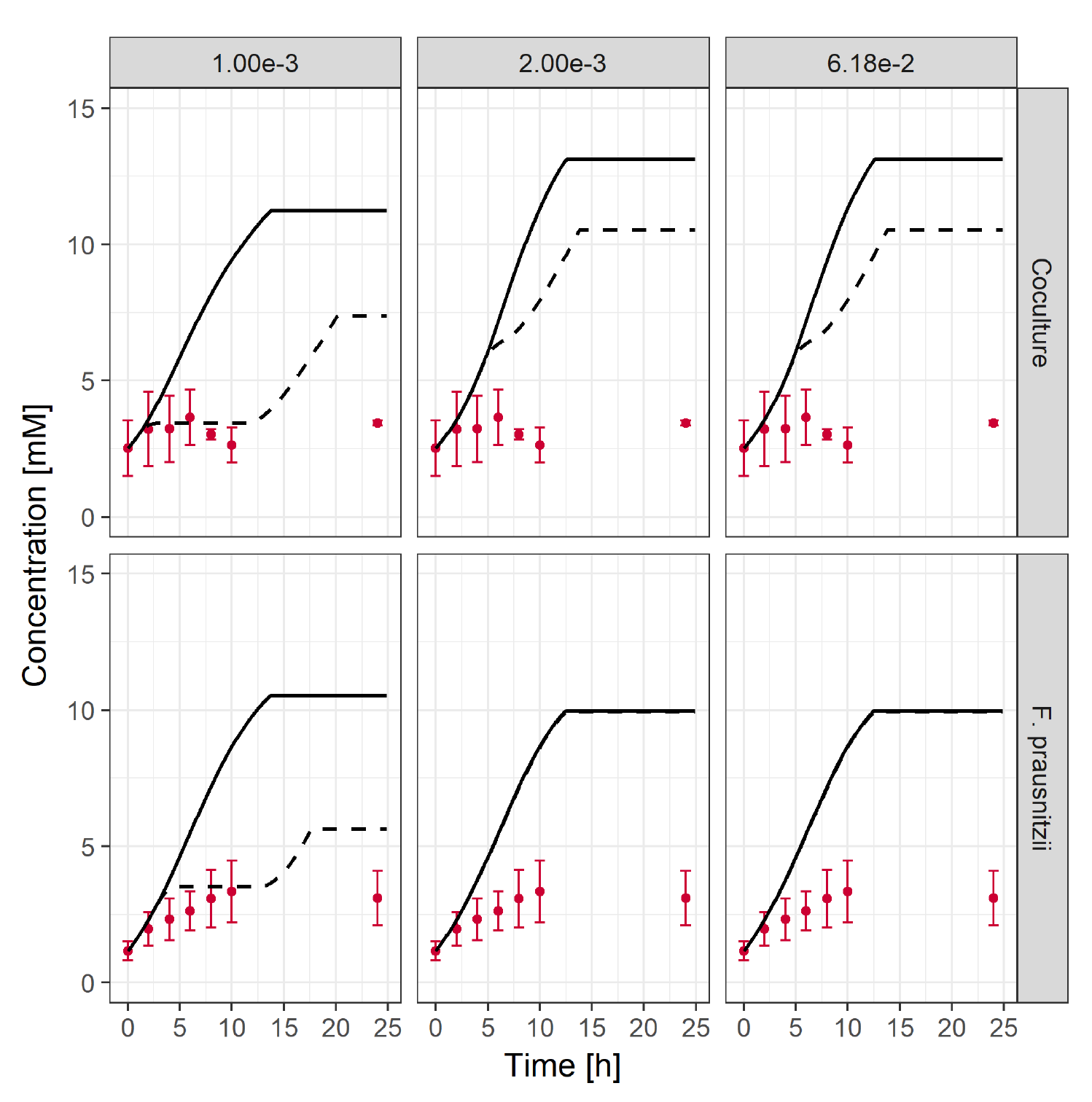
Simulated butyrate production for three starting abundances of phosphate for the low initial acetate/glucose concentration. Lowering of the phosphate concentration leads to multiple FBA solutions and difference between secondary minimization and maximization of butyrate secretion.

**Figure 5.**
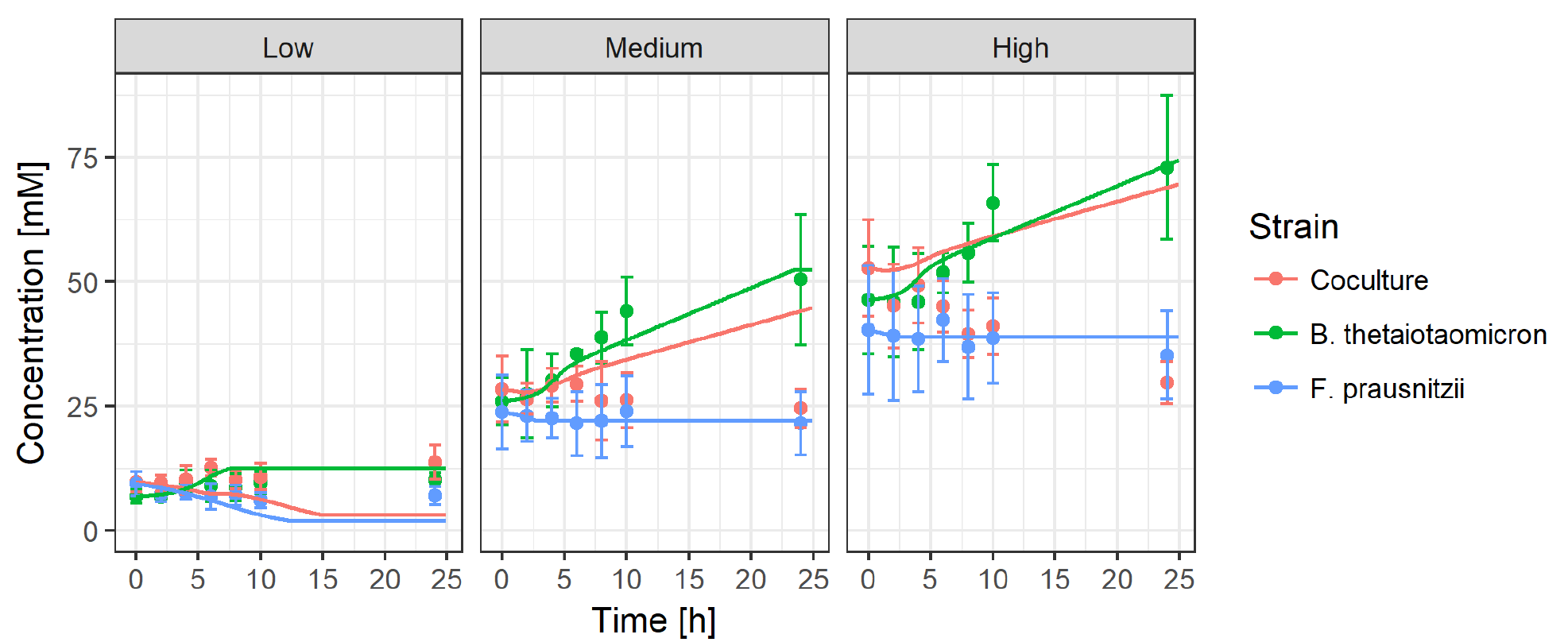
Experimental and simulated (solid curves) acetate production time courses for monocultures and coculture under the three initial glucose/acetate initial concentrations. Apparent differences demonstrated in these plots in butyrate production for *F. prausnitzii* monoculture vs. coculture and for the three initial concentrations can be tested statistically using ANOVA on summary statistics describing the curves.

**Figure 6.**
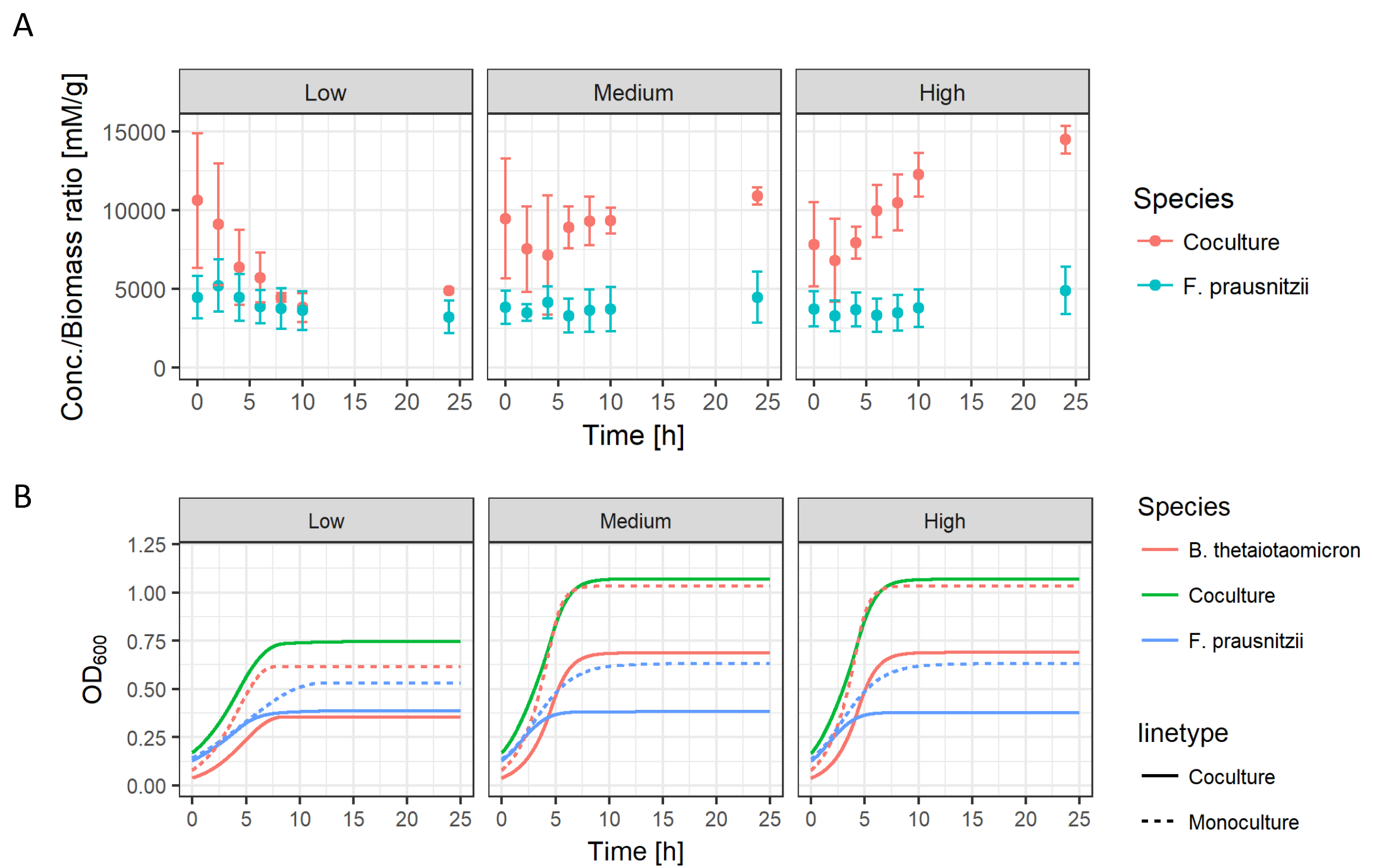
A) Butyrate concentration time profile, normalized by the simulated *F. prausnitzii* biomass, for coculture and monoculture. B) Simulated species composition.

Trends of significance in Tables 5 and 6 for the normalized curves are similar to those for the unnormalized curves, with several exceptions. The average rate of both SCFAs are significantly affected by mono vs. coculture conditions in the normalized curves. These results imply that *B. thetaiotaomicron* stimulates butyrate production in *F. prausnitzii* on a per cell basis rather than by stimulating *F. prausnitzii* biomass production. *B. thetaiotaomicron* biomass-normalized acetate production curves are not significantly changed in average rate by initial glucose/acetate levels, in contrast to the non-normalized curves. As a note of caution, it is important to stress that the normalization relative to untested predicted abundances of individual species in co-culture should be considered putative. At the same time, it could be viewed as a valuable strategy for integrating experimental and computational data towards the formulation of new hypotheses.

**Table 5.**
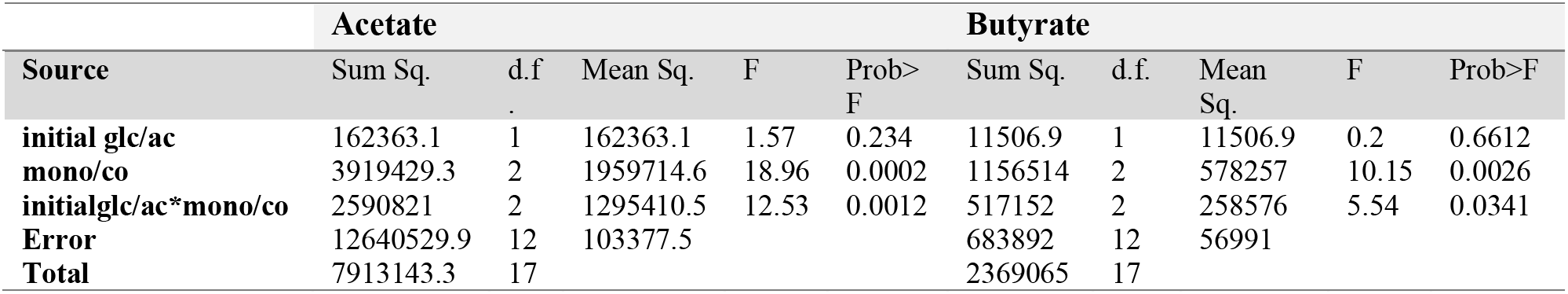
Analysis of Variance of Average Rate of Biomass-Normalized Metabolite Curves.

**Table 6.** Parameters of the Michaelis-Menten uptake functions. Glucose and acetate uptake parameters were obtained independently from the rest of the nutrients for the model of *F. prausnitzii.*

| Species | *V*<sub>max</sub>[mmol/hg] | Glc./Ac. *V*<sub>max</sub>[mmol/hg] | *K*<sub>m</sub><br>[mM] | Glc./Ac. *K*<sub>m</sub><br>[mM] |
| --- | --- | --- | --- | --- |
| *B. thetaiotaomicoron* | 11 | 11 | 3 | 3 |
| *F. prausnitzii* | 23 | 10 | 10 | 5 |

### The multiplicity of fermentation states with optimal efficiency influences SCFA time-course predictability

Figure 3 shows that COMETS accurately recapitulates the early stages of accumulation of extracellular butyrate. After 5 hours of growth, however, the picture becomes more complex. Simulation results at these times suggest that the regulation of butyrate and lactate pathways may play a major role in the final outcome of the secreted butyrate. To understand these results, the butyrate fermentation process in *F. prausnitzii* was revisited with additional computational analyses, specifically focused on alternative fermentation pathways. Three competing fermentation pathways exist in the curated metabolic network of *F. prausnitzii*. Pyruvate is fermented to one of the three products (27): (i) lactate, by D-lactate dehydrogenase (reaction ID: LDH_D_), (ii) formate, by pyruvate-formate lyase (reaction ID: PFL) or (iii) butyrate, by butyryl-CoA:acetate CoA-transferase (reaction ID: BTCOAACCOAT). Butyryl-CoA in turn is produced from acetyl-CoA (reaction ID: BTCOADH) and acetyl-CoA is converted from pyruvate by pyruvate:ferredoxin oxidoreductase (reaction ID: POR4i). The balance of metabolic flow through these three closely coupled fermentation pathways can significantly impact the production of butyrate in *F. prausnitzii,* depending on environmental conditions.

As shown in Figs. 3 and S4, and described in the methods section, COMETS, in its standard formulation, switches among these competing pathways. In particular, Figure S3 shows that *F. prausnitzii* switches from predominantly butyrate production activity at time zero to lactate secretion after 5 hours. This switch coincides with the transition from a single solution point of the FBA optimization at the early stages of the growth, to conditions where the FBA optimization algorithm has the freedom of choosing between a multitude of flux solution points, all corresponding to the same biomass growth rate. This multiplicity of equivalently efficient steady states (multiple alternative optima, also described in (37)) is best highlighted by systematically imposing, at each time point, additional features in the dFBA solution process. In particular, in analogy with flux variability analysis (38), we re-run the COMETS simulations by adding a secondary objective function at each time point. After maximizing for growth, the algorithm fixes the growth rate to the identified maximum, and subsequently searches for the solution that maximizes or minimizes the secretion flux of one of the organic acids, such as butyrate. Correspondingly, the butyrate secretion flux can be represented in the form of two curves of butyrate concentration extremes (Fig. 3). Notably, in most of the cases, the experimentally measured butyrate concentrations occupy a place between these two extremes. Nutrient limitations seem to be a strong determinant of this multiplicity of alternative optima. The system is particularly sensitive to phosphate concentration, as shown in Figs. 4 and S4, and is probably due to the strong coupling to phosphate in all butyrate producing pathways. These simulation results therefore suggest that regulation of the fermentation pathways in *F. prausnitzii* influence butyrate production under different environmental conditions.

The simulated acetate production shown in Fig. 5 tracks experimental time courses for both *B. thetaiotaomicron* and *F. prausnitzii* monoculture, particularly in the medium and high inital acetate/glucose conditions. Simulated time courses fall within error bars for average experimental observations for all time points on the curves. The early stages of the coculture simulations also closely track experiments. After 5 simulated hours, however, opposite trends in acetate concentrations appear to mirror the discrepancies with experiment in the butyrate simulations. While the experiments show depletion, the simulations result in buildup of acetate in the late stages of the simulations. This inconsistency is also potentially explainable by a metabolic switch in *F. prausnitzii*, resulting in less acetate consumption to produce butyrate.

## Discussion

We analyzed experimentally and computationally the possible effects of a symbiotic partner (*B. thetaiotaomicron*) and of environmental conditions (amount of glucose and acetate) on the biomass of the gut bacterium *F.prausnitzii*, as well as its capacity to produce butyrate. In monoculture, *F. prausnitzii* seems to continue producing butyrate even after the cells reach stationary phase (at around 10 hours), suggesting that maintenance processes keep fueling the butyrate production pathways. Another feature of the monoculture is the relatively low sensitivity of biomass and butyrate production to glucose and acetate concentrations. In contrast, biomass and butyrate seem to more strongly depend on the environment in the presence of *B. thetaiotaomicron* in the coculture experiments (Figure 3), although this difference is not statistically significant, based on ANOVA.

The possibility of metabolic cross-feeding between *F. prausnitzii* and *B. thetaiotaomicron* has been suggested in previous studies (24). In some of these studies, acetate production by *B. thetaiotaomicron* is proposed to mediate the interaction between the two bacteria, facilitating an increased butyrate production by *F. prausnitzii*. While it is likely that indeed acetate plays a key role in the interaction between the two bacteria, the results of our study suggest a more complex mode of interaction: First, the statistically significant increase in butyrate production we observe in coculture does not seem to increase monotonically with the amount of glucose and acetate, suggesting that either the two carbon sources are saturated (24), or that acetate exchange is not the only factor dominating butyrate production. Second, as demonstrated by our COMETS dFBA simulations, the stoichiometry of *F. prausnitzii* suggests that multiple alternative growth optima are possible (depending on whether or not other nutrients – most notably phosphate – are limiting in the medium). These different optima can differ substantially in their combination of fermentation products, thus making the stoichiometry-based prediction of a specific rate of butyrate production impossible. Instead, only a range of production rates can be predicted at any given time. This hypothesized degree of freedom in fermentative pathways could in principle be used by *F. prausnitzii* to modulate its metabolic activity and its butyrate production rate in response to external signals. Future studies using dFBA for studying *F. prausnitzii* could use additional constraints (e.g. total flux capacity (38) or regulatory information (38)) to refine the predictions, and to systematically test the effect of different nutrient limitations and growth media on butyrate production.

In this work, COMETS predictions were also used to estimate relative biomass amount of the two species in coculture, in the absence of experimental observations. Although the accuracy of the COMETS relative biomass estimates were not confirmed using experimental data, its ability to predict the total biomass in coculture using parameters learned from monoculture conditions lends credence to these estimates. Follow-up experiments to fully vet model accuracy in determining relative biomass in consortia would be invaluable towards building confidence in hybrid computational-experimental approaches like the one demonstrated here.

## Methods and Materials

### COMETS Simulation Configuration

The metabolic network models we use in this work for *Bacteroides thetaiotaomicron* strain VPI-5482 and *Faecalibacterium prausnitzii* strain A2-165 were published and made publicly available by Heinken *et al.* in (26) and (27). The COMETS simulation framework is implemented in Java and described in (33). R and Matlab scripts transform COMETS outputs to time-course plots. The 3D volume in these simulations contains 5 mL of isotropic medium and biomass. COMETS’ ability to model spatial differences in microbial systems is not explored in these preliminary simulations. Similarly, the dFBA settings, other than the ones mentioned below, were set at their default values as implemented in COMETS. The FBA parsimonious optimization was performed using the GUROBI optimizer, with a primary maximization of the biomass growth rate and a secondary minimization of the absolute values sum of the metabolic fluxes. The butyrate secretion analysis also included additional maximization and/or minimization of butyrate, lactate and formate uptake. Simulation run time was 24 hours, with each time step set to 0.01 hour. The death rate was set to zero.

The uptake of nutrients was modeled as a saturation Michaelis-Menten curve with two adjustable parameters, maximum uptake flux, V_max_, and the Michaelis constant K_M_. Our choice of parameters for the uptake curve was guided by values provided in the original publications of the models and as reported in the literature (39, 40). The glucose uptake in *F. prausnitzii,* for example, is governed by PTS transporter (27) with reported K_M_ values up to 8.7 mM (34). These starting values for the uptake parameters where additionally fine-tuned by fitting the single-species simulations results for the OD to the corresponding experimental curves. **Error! Reference source not found.** shows the fitting procedure for *B. thetaiotaomicron* and *F. prausnitzii* respectively. In the case of *B. thetaiotaomicron* a single value for the maximum uptake parameter was sufficient to fit the growth curves. The accepted value minimized the composite reduced chi-squared for all three growth conditions. In the case of *F. prausnitzii*, we used two values for the maximum uptake. Starting with the value for K_M_ we fitted the glucose and acetate uptake, and then performed a fine tuning for the rest of the metabolites/nutrients. Parameter values are shown in Table 6.

Metabolic activity in the *F. prausnitzii* model showed a nutrient concentration dependent shift, most sensitive to phosphate depletion, from butyrate producing pathway, to a lactate producing one, shown in Fig. S3. This shift is characterized by a single solution point of the FBA optimization sequence, both for minimized and maximized butyrate secretion, at high values of phosphate concentration, shown in Fig. S4, corresponding to the initial time in the dFBA simulation. As the substrate is depleted of the nutrients, the system obtains multiple optimal solution points, with the difference in the butyrate production depending on the secondary optimization of butyrate secretion (Fig. S4) providing a range of possible butyrate secretion rates at the later stages of the dFBA simulations. The complete set of input as well as simulations output files can be found in the supplement. COMETS is available to download at comets.bu.edu.

### ANOVA Methodology

Box’s method (36) for describing and quantifying differences in growth curves was implemented as follows. The average rate/level statistic is computed as the average of the measurements from the first time point concentration measurement subtracted from the average of the measurements from the last divided by the total time. The rate of butyrate production for each time bin was computed similarly and the average rate was subtracted from these for the set of rate deviations that define the shape statistic.

We use the Bonferonni corrected significance level of 0.0042 in this study with twelve comparisons (six per metabolite) to conservatively approximate the 0.05 significance level in single comparisons. The small number of replicates in our study (3 replicates), results in relatively low power for ANOVA tests. Insignificant results in our ANOVA analysis may therefore derive from low power, randomness or some combination of the two (41) for both the *shape* and *level* ANOVA results. Additionally, violations of homogeneous measurement covariance matrices and/or normally distributed prediction errors with zero mean could also result in pessimistically biased significance estimates. In particular, violation of homogeneous covariance matrices may negatively bias *shape* ANOVA results (42). ANOVA results for *level* have been shown to be robust to violations of this assumption (43).

We did not test our data for violations of these two conditions because tests for normality and equal variance are themselves inconclusive with small sample sizes. Given that violations of homogeneous measurement covariance matrices can negatively bias significance results in *shape* ANOVA, the insignificant results in this study should be probed further with larger sample sizes. Because *level* ANOVA results have been shown to be robust to violations of homogeneous measurement covariance matrices (43), we are confident in the finding of significant differences between monoculture and coculture metabolite production curves in *level*/average rate. As is always the case, however, follow-up studies with larger sample sizes would be advised to both test reproducibility and lend more power to ANOVA results.

### In Vitro Experimental Configuration/OD600 Analysis

Bacteria were cultured in Yeast Casitone (YC) medium, with three different concentrations of supplemented acetic acid/glucose. “Low” condition: 5.551mM acetic acid, .1% glucose. “Medium” condition: 27.754mM acetic acid, .5% glucose. “High” condition: 55.507mM acetic acid, 1% glucose. All media were adjusted to pH 6.8 before autoclaving.

Bacterial cultures were started in anaerobic conditions from glycerol stocks stored at - 80°C in 3mL of “Medium” YC medium. After overnight culture, OD600 was measured, and the cultures were diluted to the nominal OD starting points in 5mL of the three different YC formulations.

The following cultures were started with the initial OD600 values as noted:

OD600 ~.02 *B. thetaiotamicron* monoculture

OD600 ~.08 *F. prausnitzii* monoculture

OD600 ~.02 *B. thetaiotamicron* AND OD600 ~.08 *F. prausnitzii* coculture

A baseline 200μL aliquot was taken from each culture, measured by OD600, and stored at −80 for later MS analysis. Subsequent 200μL aliquots were collected and measured by OD600 at 2, 4, 6, 8, 10 and 24 hours and stored for later analysis. The above procedure was repeated in triplicate, yielding three observations per time point.

### MSMS Analysis

A flow injection analysis electrospray ionization mass spectrometry (FIA ESI MSMS) method was used for quantitative detection of short chain fatty acids (SCFA). Acetic, propionic, butyric and succinic acid were derivatized with 3-nitrophenylhydrazine in the presence of N-(3-dimethylaminopropyl)-N’-ethylcarbodiimide and pyridine and detected by a mass spectrometer as a 3-nitrophenylhydrozones in MRM (multiple reaction monitoring) MSMS mode, as described by J. Han *et al.* (44). To increase precision and robustness of the method, 3- methylbutyric-2-2-d2 acid, acetic acid-2-13C and propionic acid −1-13C were used as an internal standards. Quantitation was done by external standards calibration, where instrument response for the analyte was measured as a ratio between analyte’s and internal standard’s peak areas. The FIA technique did not utilize LC column but rather a direct injection of the sample into an ESI probe of the mass spectrometer and this decreased time of analysis per sample to two minutes. The FIA ESI MSMS for the detection of SCFA is sensitive with the limit of detection for acetic, propionic, butyric and succinic acids at 4, 3, 0.6 and 1.4μM respectively. The accuracy of the method was between 98-102%.

### Reagents

LC MS grade acetonitrile and water were purchased from VWR (Radnor, PA, USA). N-(3-dimethylaminopropyl)-N’-ethylcarbodiimide HCl, 3-nitrophenylhydrazine HCl, pyridine, acetic acid, propionic acid, butyric acid, succinic acid, acetic acid 13C, and propionic acid 13C were purchased from Sigma-Aldrich (St Luis, MO, USA). 3-methylbutyric-2-2-d2 acid was purchased from CDN Isotopes (Quebec, CN).

### FIA MS/MS system

An Agilent infinity capillary LC pump with micro-autosampler and thermostat (Agilent Technologies, Santa Clara, CA, USA) coupled to AB Sciex 4000 Q-TRAP triple-quadrupole mass spectrometer (AB Sciex, Concord, Ontario, CN) was used for the analysis. The flow solvent - five percent water and ninety five percent acetonitrile was delivered to a mass spectrometer ESI probe at the rate of 350μL/min. Samples for flow injection analysis were derivatized on the Agilent polypropylene 96 well plate and injected into mass spectrometer with injection volume of 40μL. Following conditions for the AB Sciex Q-TRAP 4000 were used for analysis: source temperature 400°C, source gas 40L/min, curtain gas 10L/min, ESI capillary voltage was set at –4500 volts. Data were acquired in negative polarity multiple reactions monitoring (MRM) mode for the MRM transitions specified in the Table S1.

## Data Availability

Data files and scripts used to generate the figures presented in this paper can be found in a zipped directory (Yu_etal_Data_and_Scripts.zip) downloadable at https://github.com/segrelab/Fprau-Btheta-2018. This directory contains the experimental data and script for statistical analysis and for generating the figures (DATA_AND_FIGURE_SCRIPTS subdirectory), and COMETS input and output files (SIMULATIONS_INPUTS_AND_OUTPUTS subditrectory). The *in silico* experiments were generated using COMETS v.2.5.8, which is freely available at http://comets.bu.edu.

## Acknowledgements

DS and ID acknowledge funding from the Defense Advanced Research Projects Agency (Purchase Request No. HR0011515303, Contract No. HR0011-15-C-0091), the NIH (5R01DE024468, R01GM121950 and Sub_P30DK036836_P&F), and the Boston University Interdisciplinary Biomedical Research Office. This material is based upon work supported by the Assistant Secretary of Defense for Research and Engineering under Air Force Contract No. FA8702-15-D-0001. Any opinions, findings, conclusions or recommendations expressed in this material are those of the author(s) and do not necessarily reflect the views of the Assistant Secretary of Defense for Research and Engineering.

## Supplemental table legend

Table S1. AB Sciex 4000-TRAP parameters used for detection of SCFA Q1 – m/z of the analyte ion detected on the 1rst quadrupole.Q3 – m/z of the anatyte’s fragment ion detected on the third quadrupole. DP-declustering potential, CE – collision energy. Analytes in red are internal standards.

## Supplemental figure legends

**Figure S1.** Simulated glucose concentration.

**Figure S2.** Biomass normalized acetate concentration.

**Figure S3.** Simulated fluxes of key reactions in the butyrate fermentation pathway in *F. prausnitzii,* for high glucose concentration, at time zero and 5 hours.

**Figure S4.** Simulated butyrate secretion fluxes in *F. prausnitzii*, under secondary maximization/minimization of butyrate or lactate production, as a function of phosphate concentration, in low glucose conditions.

